# Bnip3 expression is strongly associated with reelin-positive entorhinal cortex layer II neurons

**DOI:** 10.1101/2023.10.19.562876

**Authors:** Stig W. Omholt, Raissa Lejneva, Maria Jose Lagartos Donate, Evandro Fei Fang, Asgeir Kobro-Flatmoen

## Abstract

In layer II of the entorhinal cortex, the principal neurons that project to the dentate gyrus and the CA3/2 hippocampal fields express the large glycoprotein reelin (Re+ ECLII-neurons). In rodents, neurons located at the dorsal extreme of the entorhinal cortex that thus border the rhinal fissure, express the highest levels, and the expression gradually decreases at levels successively further away from the rhinal fissure. Here we test two predictions following from the hypothesis that reelin expression is strongly correlated with neuronal metabolic rate. Since mitochondrial turnover rate serves as a proxy for energy expenditure, we predicted that the expression of the canonical promitophagic BCL2 and adenovirus E1B 19-kDa-interacting protein 3 (Bnip3) would be upregulated in Re+ ECLII-neurons, and that the degree of upregulation would strongly correlate with the expression level of reelin in these neurons. We confirm both predictions, which implies that the energy requirement of Re+ ECLII-neurons is generally high, and that there is a systematic decrease in metabolic rate in these neurons as one moves successively away from the rhinal fissure. We tentatively suggest that the reasons for the high energy requirement of these neurons are their high rate of synaptic transmission and the high frequency by which they remold their synaptic contacts. This implies that the systematic variation in energy requirement of the neurons manifesting the observed reelin gradient ties in with the level of spatial and temporal detail by which they encode information about the external environment.

## Introduction

The entorhinal cortex (EC) is located deep in the medial temporal lobe where it processes an enormous amount of information, mediating a large majority of the information flow between the neocortex and the hippocampus^1^. Functionally, EC plays a crucial role in the formation, consolidation, and retrieval of memories and spatiotemporal representations^2^. The definition of EC is anchored to the fact that it projects to the hippocampal dentate gyrus, aligned with specific cytoarchitectonic features, and in rodents, two main subdivisions are easily recognized, namely the medial EC (MEC) and the lateral EC (LEC) ^3^. These subdivisions have their counterparts in primates, including humans, in which the anteriolateral EC approximately corresponds to LEC, and the posteromedial EC approximately corresponds to MEC^3-6^. Since, in particular, the population of large clustered projection neurons in the lateral or anterolateral EC layer II (LII) shows clear signs of degeneration from the earliest stages of AD^7-10^, this domain of ECLII is clearly involved in the very early etiology of the disease^11^. Therefore, a better understanding of native ECLII physiology is certainly of biomedical interest.

In the adult brain, the large glycoprotein reelin functions as a potent inducer of synaptic spine-growth^12,13^ and plasticity^14^, and promotes long-term potentiation^15^. Reelin is mainly expressed in a subset of interneurons found throughout the cortex. However, in ECLII, the subset of principal neurons that projects into the dentate gyrus and the CA3/2 hippocampal fields expresses a high level of reelin (Re+ ECLII-neurons). The Re+ ECLII-neurons express reelin in a strikingly graded pattern. In rodents, neurons located at the dorsal extreme of EC, that is, at the border of the rhinal fissure, express the highest levels, and the expression gradually decreases toward the ventral extreme^16^. Correcting for the different gross orientation of EC between different orders of mammals, the same pattern holds true for all mammals examined, including humans^3^.

The morphology of Re+ ECLII-neurons strongly suggests that they are heavily involved in synaptic transmission. Most Re+ ECLII-neurons have a stellate (MEC) or fan (LEC) morphology with imposing dendritic trees. A single Re+ ECLII-neuron in a rat has a densely spinous dendritic tree whose distal 360 degree diametric perimeter can span upwards of 900 μm^17,18^. To appreciate the size of this span, one may consider another cell type with very large dendritic trees, namely hippocampal pyramidal neurons. These neurons have apical dendritic arbors estimated to be four-to-nine times larger and basal dendritic arbors approximately three times larger than that of a pyramidal neocortical neuron^19^. However, the apical (and largest) dendritic arbor of a large pyramidal hippocampal neuron, measured at its distal perimeter^19,20^, is still only about half that of a Re+ ECLII-neuron. Likely facilitated by their formidable dendritic trees, *a single* Re+ LECLII-neuron is capable of collecting direct input from at least five separate cortical areas, in addition to multiple sources of subcortical modulatory input^11^. Furthermore, these neurons give rise to particularly extensive axonal arborizations both in their downstream hippocampal target zones and locally in EC^21^. Since most of the brain energy is spent on synaptic transmission^22^, the energy requirement of these neurons is likely very high. In addition, the presence of high levels of reelin implicates a particularly high need for remolding synaptic contacts, which further enhances their energy requirement.

The above considerations led us to hypothesize that Re+ ECLII-neurons in general have a particularly high metabolic rate and that the rate is highest in those located closest to the rhinal fissure. Since mitochondria provide the lion’s share of brain energy^22^, mitochondrial turnover rate appears to be a superb proxy of metabolic rate under normal physiological conditions as it probably strongly correlates with both the number of mitochondria and the metabolic demand on each mitochondrion, i.e., it correlates with the frequency with which impaired mitochondria arise. Thus, our hypothesis would be supported by establishing that the mitochondrial turnover rate in Re+ ECLII-neurons is strongly associated with their reelin expression.

The main function of BCL2 and adenovirus E1B 19-kDa-interacting protein 3 (Bnip3) is to induce mitophagy^23-25^ and the mRNA precursor of Bnip3 is especially enriched in the EC of adult rats^26^. Compromised mitophagy is associated with age-related diseases, including AD^27,28^, and uncovering putatively differentially expressed mitophagic proteins such as Bnip3 is, in addition to its relevance for metabolic rates, also relevant for our understanding of brain region-specific vulnerabilities. These considerations led us to assay the relative amount of Bnip3 in the brain of rats. We found that although nearly all areas contained none or very low levels of Bnip3, which is in line with previous findings^29-32^, ECLII constitutes a striking exception. Double-immunofluorescence labeling revealed that nearly every Re+ ECLII neuron expresses Bnip3 and that the expression level of Bnip3 aligns with the reelin gradient. Since there appears to be a consonance between the reelin and Bnip3 gradients, how Re+ ECLII-neurons connect to the hippocampal formation^3^, and the resolution of encoded environmental representations, we suggest that the systematic variation in energy requirements and reelin expression in Re+ ECLII-neurons may possibly reflect the frequency by which they need to update information about the external environment.

## Methods

For this study we used eight outbred Wistar Han rats purchased from Charles River (France). We divided the animals into four age-groups, 3-months, 6-months, 12-months, and 18-months, each containing two rats. All procedures were approved by the Norwegian Animal Research Authority, and follow the European Convention for the Protection of Vertebrate Animals used for Experimental and Other Scientific Purposes. We kept the animals on a 12-h light/dark cycle under standard laboratory conditions (19–22 °C, 50–60% humidity), with access to food and water ad libitum. The human case was from archival material obtained from King’s College London Biobank, and its use was approved by The Regional Committees for Medical and Health Research Ethics, Norway (permit 82685)

### Processing and immunohistochemistry

We fully anaesthetised the animals by placing them in a chamber containing 5% isoflurane gas (Baxter AS, Oslo, Norway), followed by an intraperitoneal injection of pentobarbital (0.1 ml per 100 g; SANIVO PHARMA AS, Oslo, Norway). Subsequently we carried out transcardial perfusions with Ringer’s solution at room temperature (in mM: NaCl, 145; KCl, 3.35; NaHCO3, 2.38; pH ∼6.9) immediately followed by circulation of 4% freshly depolymerised paraformaldehyde (PFA; Merck Life Science AS/Sigma Aldrich Norway AS, Oslo, Norway) in a 125 mM phosphate buffer (PB) solution (VWR International, Pennsylvania, USA), pH 7.4. The brains were extracted and post-fixed overnight in PFA at 4°C, and then placed in a freeze protective solution (dimethyl sulfoxide in 125 mM PB with 20% glycerol, Merck Life Science AS/Sigma Aldrich Norway AS, Oslo, Norway) until sectioning. We sectioned the brains coronally at 40 μm into six series, using a freezing sledge microtome.

Immunohistochemistry was done on free-floating sections. To improve access to the antigen, the sections were subjected to gentle heat induced antigen retrieval by placement in PB at 60°C for two hours. <u>For chromogenic immunoenzyme-labelling:</u> To avoid unspecific labelling the sections were first incubated for 2 × 10 minutes in a solution of PB with 3% hydrogen peroxide (Merck Life Science AS/Sigma Aldrich Norway AS, Oslo, Norway), and next incubated for 1 hour in a solution of PB with 5% normal goat serum (Abcam Cat# ab7481, RRID: AB_2716553). Subsequently, free-floating sections were incubated with the primary antibody, rabbit anti-BNIP3 (BioNordika, Oslo, Norway, Cat# CST-3769S, RRID:AB_2259284) in a 1:500 dilution in PB containing 0.4% Saponin (VWR International AS, Bergen, Norway) and 5% normal goat serum, for 48 hours. We also tested a primary incubation time of 20 hours (i.e. overnight), which gave slightly less overall signal intensity. We opted for 48 hours to make sure we did not miss any potentially weakly positive neurons or neuropil. Sections were then rinsed for 3 × 5 minutes in Tris-buffered saline (50 mM Tris, 150 mM NaCl, pH 8.0) containing 0.4% Saponin (TBS-Sap), and then incubated with biotinylated secondary antibody goat anti-rabbit (1:400; Sigma-Aldrich Cat#B8895, RRID: AB_258649) in TBS-Sap, for 90 minutes at room temperature. Subsequently, the sections were rinsed in TBS-Sap, and then incubated in Avidin-Biotin complex (ABC, Vector Laboratories Cat# PK-4000, RRID: AB_2336818) for 90 minutes at room temperature. The sections were then rinsed again, and then incubated in a solution containing 0.67% 3,3’-Diaminobenzidine (Sigma/Merck, Cat# D5905) and 0.024% hydrogen peroxide for 4 minutes. We safeguarded against potentially subtle differences that can arise from small variations in the time of incubation or the solution with the chromogen by incubating sections from different age groups in parallel, using the same solution of Diaminobenzidine. <u>For immunofluorescence labelling:</u> All tissue was first subjected to Heat Induced Antigen Retrieval, by immersion in PB at 60°C for 2 hours, and then blocking against unspecific labelling was done by placing the tissue in 5% goat serum in PB for 1 hour. For reelin and Bnip3, we incubated the sections overnight with primary antibodies (mouse anti-reelin G10 clone (1:1000), Cat#MAB5364, RRID:AB_2179313 and rabbit anti-BNIP3 (1:500), BioNordika, Oslo, Norway, Cat# CST-3769S, RRID:AB_2259284) in PB solution containing 0.4% Saponin (VWR, Cat# 27534.187) and with 5% goat serum at 4 degrees Celsius. The sections were subsequently incubated for two hours with secondary antibodies, including Alexa 546 Goat anti-Mouse (Cat#A11003, RRID: AB_10562732) and Alexa 488 Goat anti-Rabbit (Cat# A11008, RRID: AB_10563748), in a solution containing 0.4% Saponin and 5% goat serum at room temperature. For Bnip3 and tyrosine hydroxylase (TH) double-immunolabeling, we used the same approach as above for Bnip3, while for TH we used mouse anti-tyrosine hydroxylase (Millipore, Cat# MAB318). Processed tissue was mounted on Superfrost™ glass slides (Thermo Fisher Scientific) from a solution of 50 mM tris(hydroxymethyl)aminomethane (Millipore, Cat# 1.08382.1000) with hydrochloric acid, at pH 7.6, and then left to dry overnight before being coverslipped using a mixture of Xylene (VWR International AS, Bergen, Norway) and entellan (Merck KGaA, Cat# 1.07960.0500).

The human tissue consisted of a single 6 μm section of EC mounted on glass. Prior to immunohistochemical procedures, the tissue was deparaffinised by immersion in xylene for 5 minutes x 3 times. The tissue was then rehydrated by being dipped 10 times in water containing decreasing amounts of ethanol (100%, 90%, 70%, 50%, 0%). We blocked against unspecific antigens by applying DAKO Real blocking solution (DAKO, Cat# S2023) for 10 minutes, followed by application of normal goat serum (same as above) for 1 hour. To detect Bnip3 we used the same antibody as for the rat tissue (see above) in the same solution as listed above, and incubated the tissue overnight. The secondary antibody and visualization-step was also as that described for the rat tissue.

### Imaging and quantification

All tissue sections were scanned on an automated Zeiss Axio Scan.Z1 system with a 20x objective (Plan-Apochromat 20x/0.8 M27), using identical settings. Analyses and quantifications were done in Zen software (2.6, Blue Ed.). For our quantifications, we used a series of sections covering the rostrocaudal extent of EC from each of three animals, and double-immunolabeled against reelin and Bnip3. While visualizing only the channel used to scan the reelin signal (546-channel), we selected cell profiles using the Zen Circle Tool, taking care to obtain good coverage also of the rostrocaudal axis, but avoiding the part at the ventral extreme (i.e. the part furthest away from the rhinal fissure) since the signal here is very weak relative to background. After this selection, we switched to the channel for the Bnip3 signal (488-channel) and identified each circle with a clearly positive signal. By this we obtained the ratio between Bnip3-positive neurons and reelin-positive neurons, i.e. 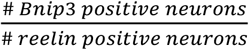.

### Orientation of sections

To aid visualisation of the position of the sections shown in the Results, we provide a cartoon (Fig. 1).

**Figure 1.**
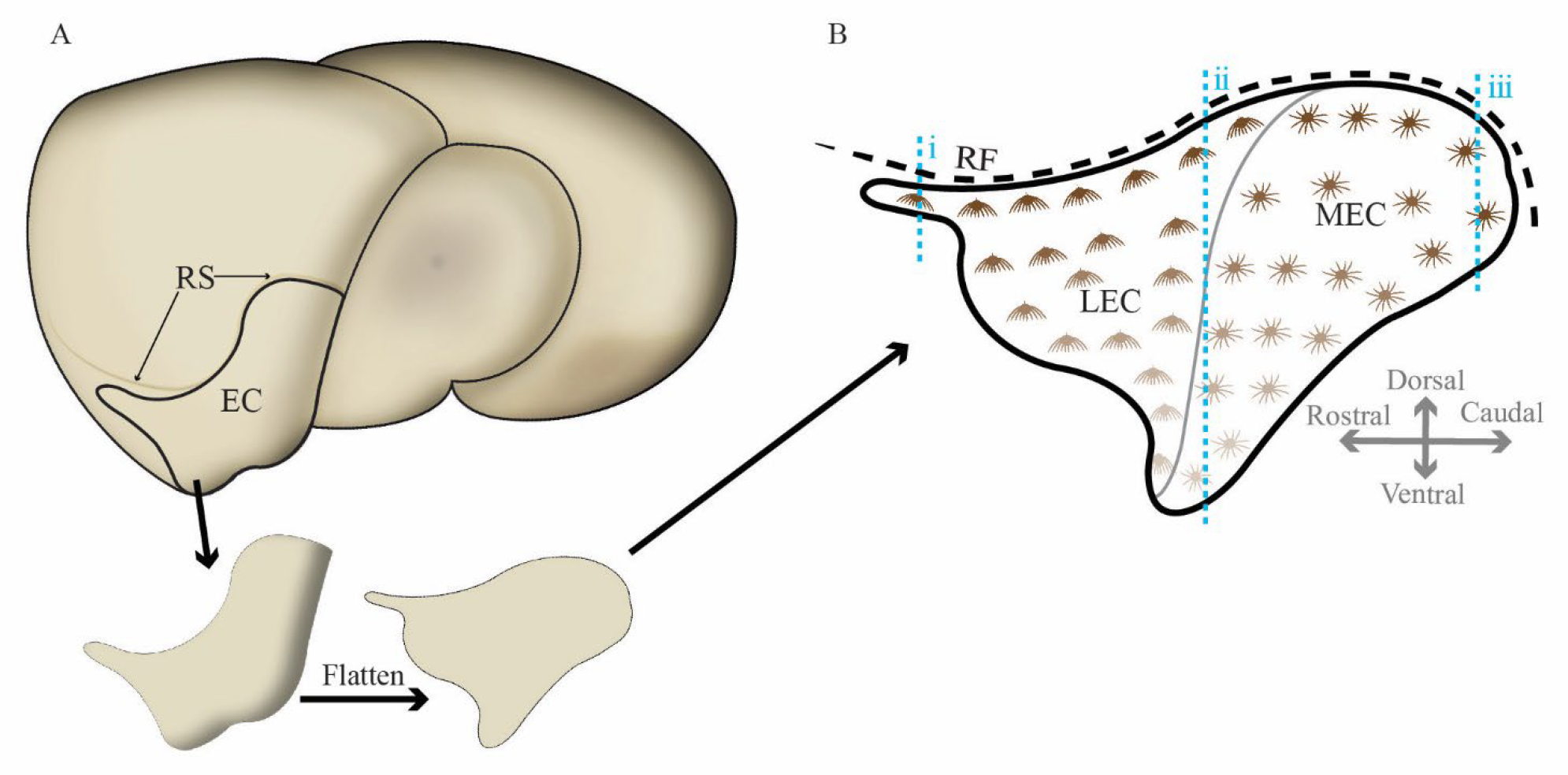
Guide to orientation of sections. (a) Cartoon of rat brain seen from a laterocaudal perspective, with the situation of the entorhinal cortex (EC) indicated, and an illustration of a virtual resection and compression of EC into a flatmap. (b) Matching flatmap of EC layer II with reelin-expressing fan cells illustrated for the lateral entorhinal cortex (LEC) and reelin-expressing stellate cells illustrated for the medial entorhinal cortex (MEC). The decrease in color intensity for the cells indicate the decrease in reelin-expression as one moves successively away from the rhinal fissure (RF). Blue dashed lines (i-iii) indicate the levels that the coronal sections in figure 3 (a-c, respectively) are taken from.

## Results

### Specificity of the Bnip3 antibody

To test the specificity of the Bnip3 antibody, we immunolabeled sections containing EC from different age groups and omitted the primary antibody on one section per run. We found no signs of background labeling in our setup (Fig. 2 and Fig 3 b).

**Figure 2.**
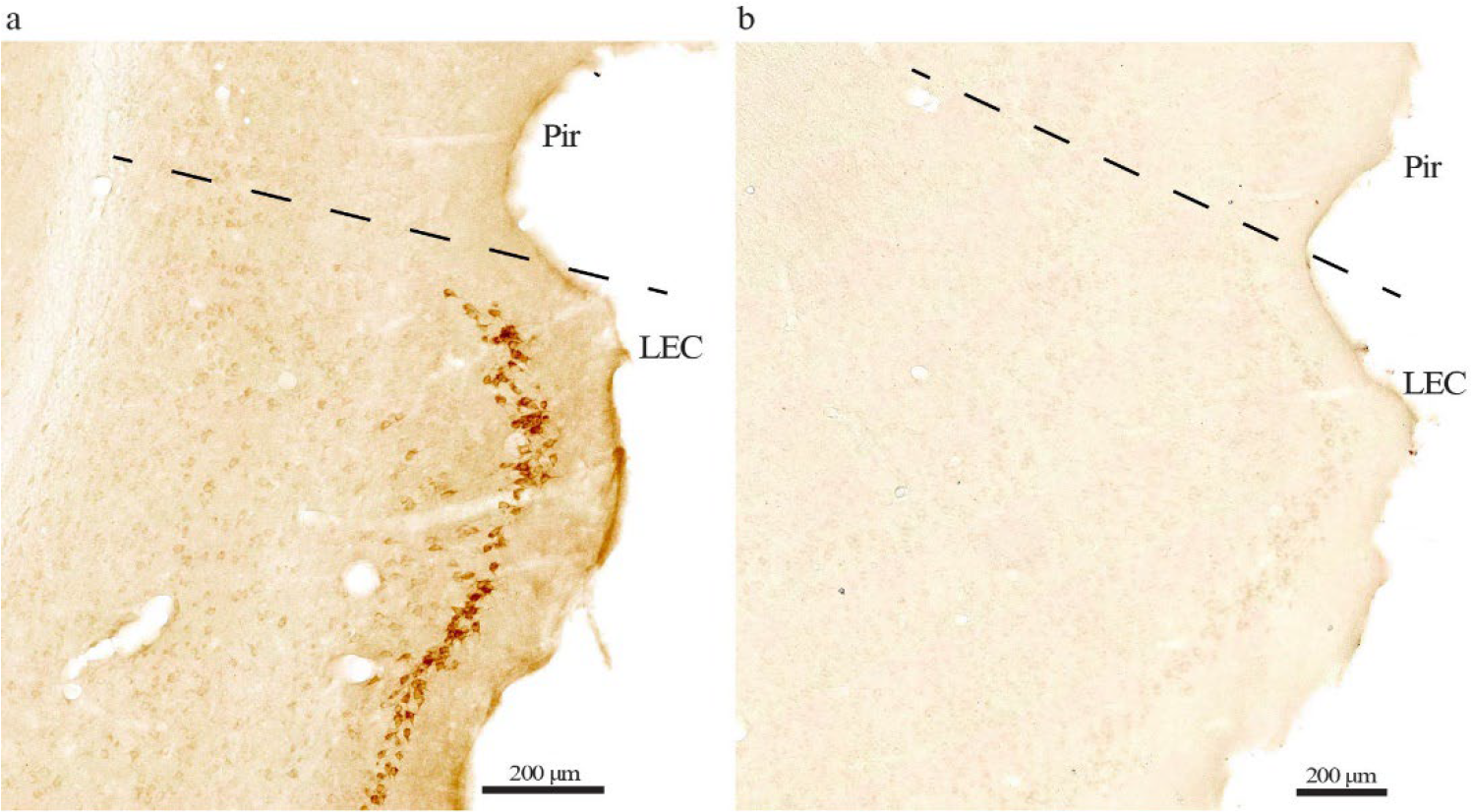
No background labeling evident in our setup. (a) Immunolabeling with the Bnip3 antibody reveals a high level of signal in cells in LEC layer II of 18-month-old Wistar rats. (b) Running the same experiment without the Bnip3 antibody while keeping all else equal, shows a complete absence of signal. The same is the case across the age groups. Scale bars indicated on each figure. Dashed line marks border between LEC and the piriform cortex (Pir).

**Figure 3.**
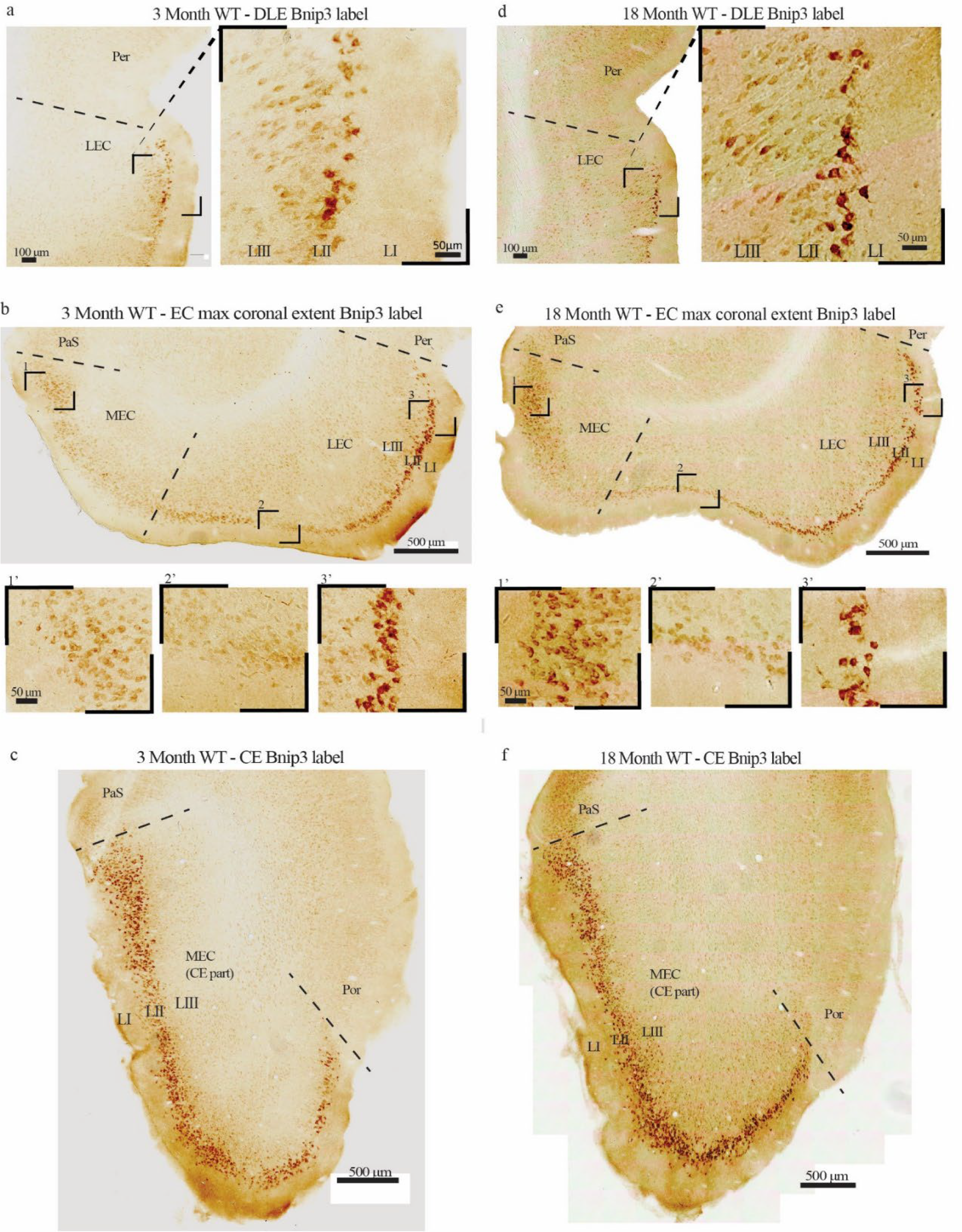
A major subset of ECLII neurons expresses Bnip3. (a) Example of the dorsolateral EC (DLE, i.e. the anterior tip of LEC), in which LII-neurons are particularly strongly stained. (b) The expression of Bnip3 gradually decrease at positions successively further away from the rhinal fissure, irrespectively of whether one is located in LEC or MEC (see figure 1 as a guide to how the sections relate to each other across EC and between LEC vs MEC). (c) Example at the caudal extreme (CE) of EC where only the part of MEC located close to the rhinal fissure is present in coronal sections. Abbreviations: Per = Perirhinal cortex, PaS = Parasubiculum, Por = Postrhinal cortex. Scale bars are indicated for each figure.

### Bnip3 expression in the entorhinal cortex

We found that the expression of Bnip3 is or undetectable or near undetectable in the cell somata of most parts of the forebrain. This is in line with previous reports^29-32^ and the finding that pathological conditions such as hypoxia or ischemia are required to trigger increased expression^30-32^. However, in stark contrast to this general finding, a major subset of neurons in the part of EC LII that is located close to the rhinal fissure (i.e., dorsally) expressed high levels of Bnip3 in all age groups (Fig. 3). Since the only difference between young and old rats appears to be a modest overall increase in staining intensity with age, the description to follow holds true for both age groups. The expression of Bnip3 in both LEC and MEC manifests a striking gradient (Fig. 3 b and c). While the expression is high in LII-neurons located close to the rhinal fissure, the expression level gradually decreases as one moves successively further away from the rhinal fissure, such that the level in the most ventrally located LII-neurons is almost below the detection limit.

### Bnip3-expression is restricted to the reelin-positive ECLII-population

Assuming that the level of Bnip3 expression serves as a proxy of the neuronal mitochondrial turnover rate, and that the turnover rate is positively correlated with the neuronal metabolic rate, it follows from the data (Fig. 3) that the overall neuronal metabolic rate is substantially higher in EC LII than in other layers of the EC, as well as in most other parts of the rat brain (see below). And the data clearly suggest that energy expenditure is highest in neurons located close to the rhinal fissure.

Since the strength of the Bnip3 signal clearly appeared to align with the previously reported reelin expression gradient of Re+ ECLII-neurons^16^, we double-immunolabeled against reelin and Bnip3. The results (Fig. 4) clearly suggest that high Bnip3 expression is restricted to the population of Re+ ECLII-neurons.

**Figure 4.**
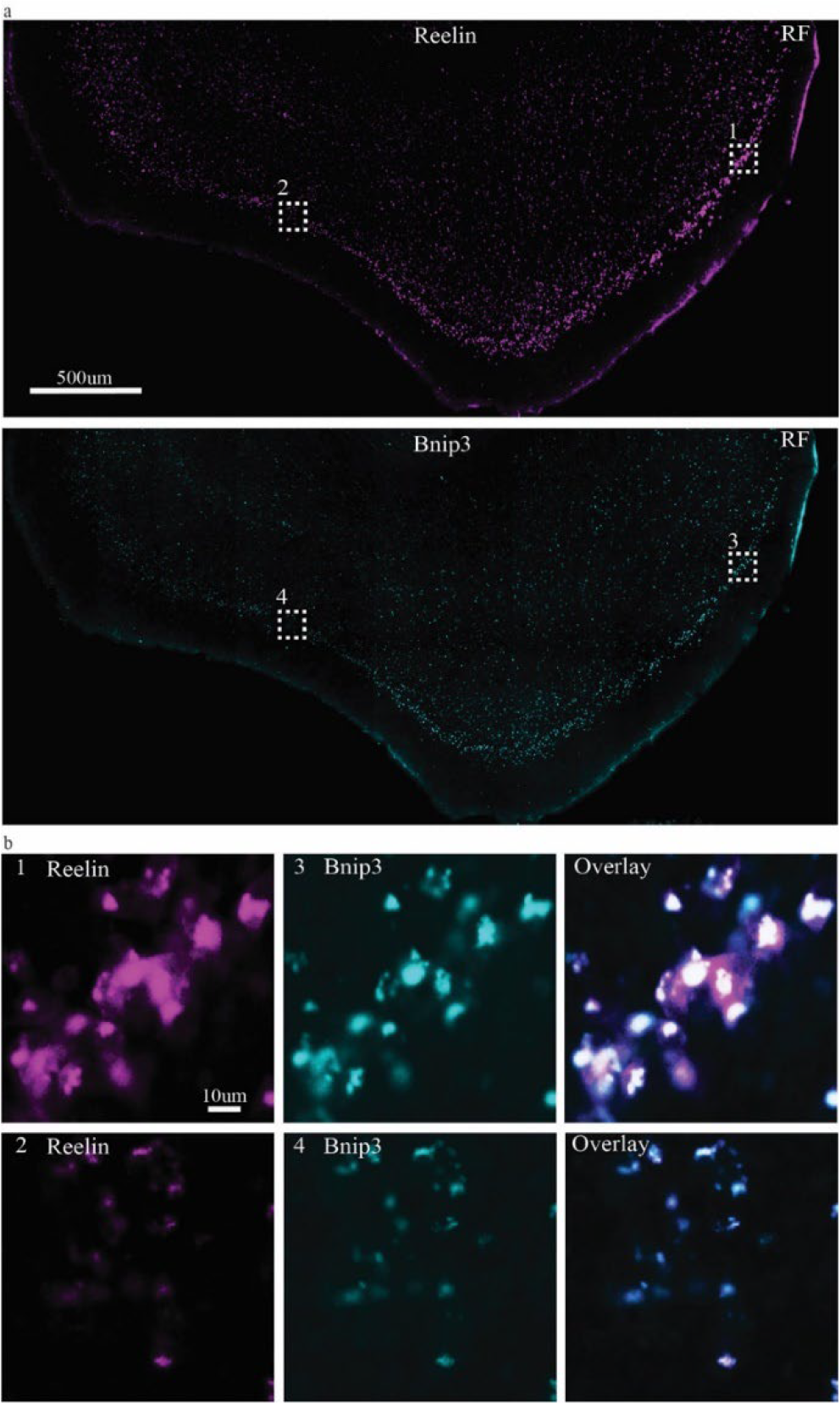
Double-immunolabeling against reelin and Bnip3. (a) Top panel shows labeling for reelin, bottom panel shows labeling for Bnip3. (b) Higher powered insets (numbers correspond to those in (a)) reveal that Bnip3 follows the same gradient as reelin, thus being strong towards the rhinal fissure (RF) and increasingly less so as one moves successively further away from the rhinal fissure. Further, it is apparent from these images that Bnip3 is confined to the population of Re+ ECLII-neurons. Scalebar for (a) applies to both images; scalebar for (b1) applies to all six images.

We therefore took one series of sections from each of three animals and quantified the degree of colocalization between Bnip3 and reelin throughout ECLII. Applying stringent criteria for what qualifies as a positive Bnip3 signal, we found that Bnip3 expression was restricted to Re+ ECLII-neurons, and that approximately 90% of Re+ ECLII-neurons were positive for Bnip3 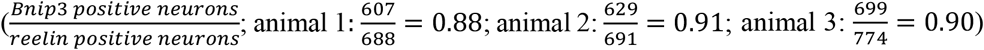. This is probably a conservative estimate, and we expect that more extensive analyses are likely to find that the degree of overlap is even higher. Given that the Bnip3 expression level serves as a proxy for neuronal metabolic rate, it appears that the data support our prediction (see Introduction) that the observed reelin-gradient in ECLII reflects the overall energy expenditure of the neurons along this gradient.

### Prediction of Bnip3 expression pattern in human EC LII

While we found Bnip3-expression also in human ECLII neurons (Fig. 5), we do not currently have access to complete human EC specimens and therefore cannot assess whether all our results obtained in Wistar rats extend to humans. However, as in rats, reelin is expressed in human principal ECLII neurons following the same topographical gradient^3^. We therefore consider it likely that a close overlap between reelin expression and Bnip3 will prove to be the case also in humans.

**Figure 5.**
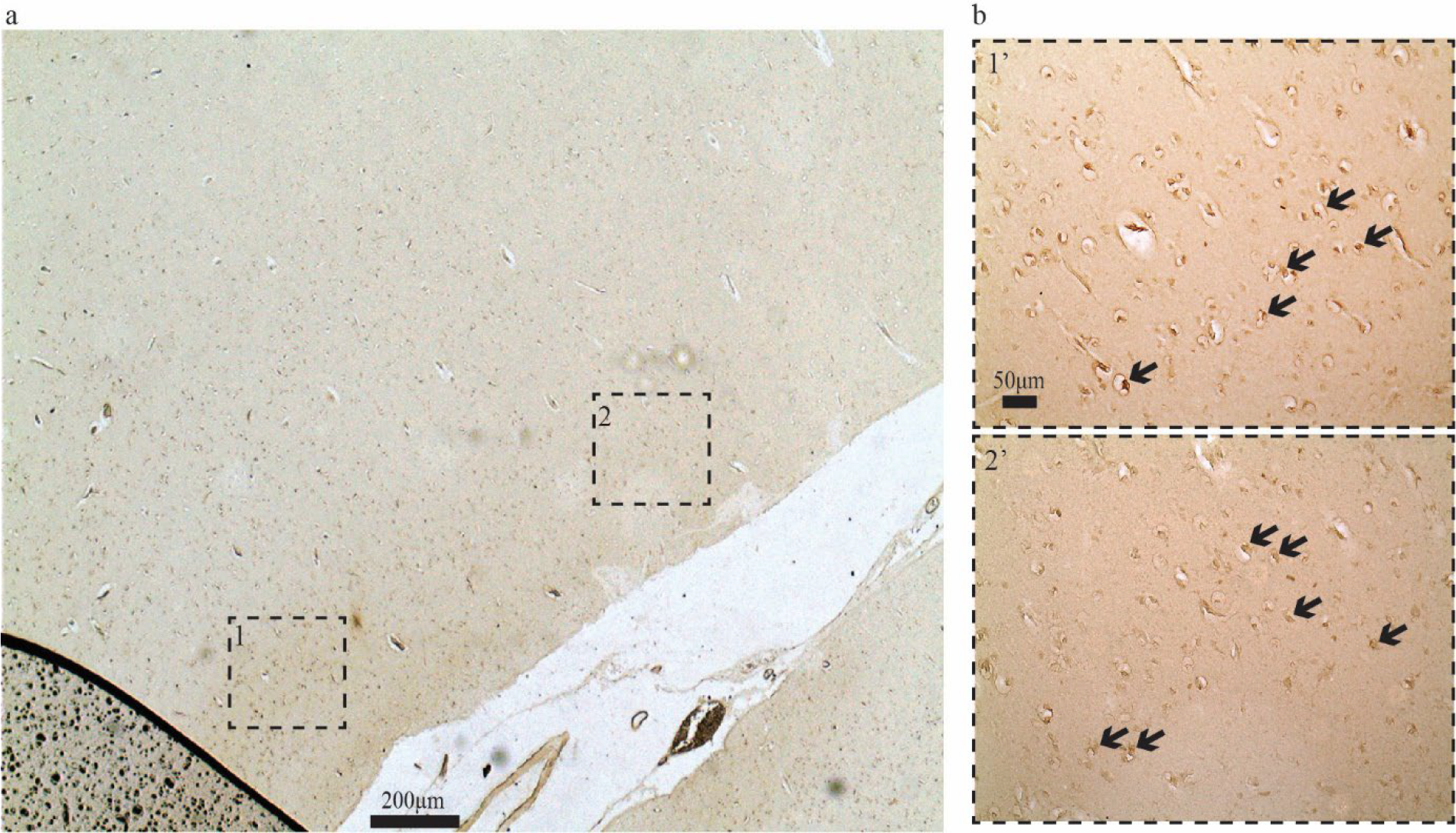
Bnip3-expression in human ECLII neurons. (a) Immunolabeling against Bnip3 in EC of a human subject appears to reveal a signal that is (b) most notable in LII (the insets in (b) match those in (a)). Arrows in (b) indicate examples of what are likely positive neurons. Scale bars are indicated in figures, (b 1’) also applies to (b 2’).

### Other neurons expressing elevated levels of Bnip3 in rats

We then used Bnip3 as a probe to identify neurons or neuronal clusters outside EC that putatively have higher than average mitochondrial activity (Fig. 6). LII-neurons of the piriform cortex and occasional neurons in the preoptic nucleus display moderate and high levels of Bnip3 expression, respectively (Fig. 6 a). In the hippocampal formation, the expression of Bnip3 is generally low or absent, except for field CA3, where a substantial number of neurons in the pyramidal layer express moderate levels. Notable levels of Bnip3-expression is also found in the outer 2/3ds of the molecular layer of the dentate gyrus (DG) and in stratum lacunosum moleculare of CA3/2, which contains the Re+ ECLII neuron-terminals; we also observe that there is a tendency for more intense labeling in the free (lower) blade of DG (Fig. 6 b). Furthermore, in parts of the visual cortex (Fig. 6 c), the retrosplenial, somatosensory and motor cortices (Fig. 6 d), and the frontal pole (Fig. 6 e) we find subsets of neurons where Bnip3 expression is clearly above background, sometimes reaching moderate levels.

**Figure 6.**
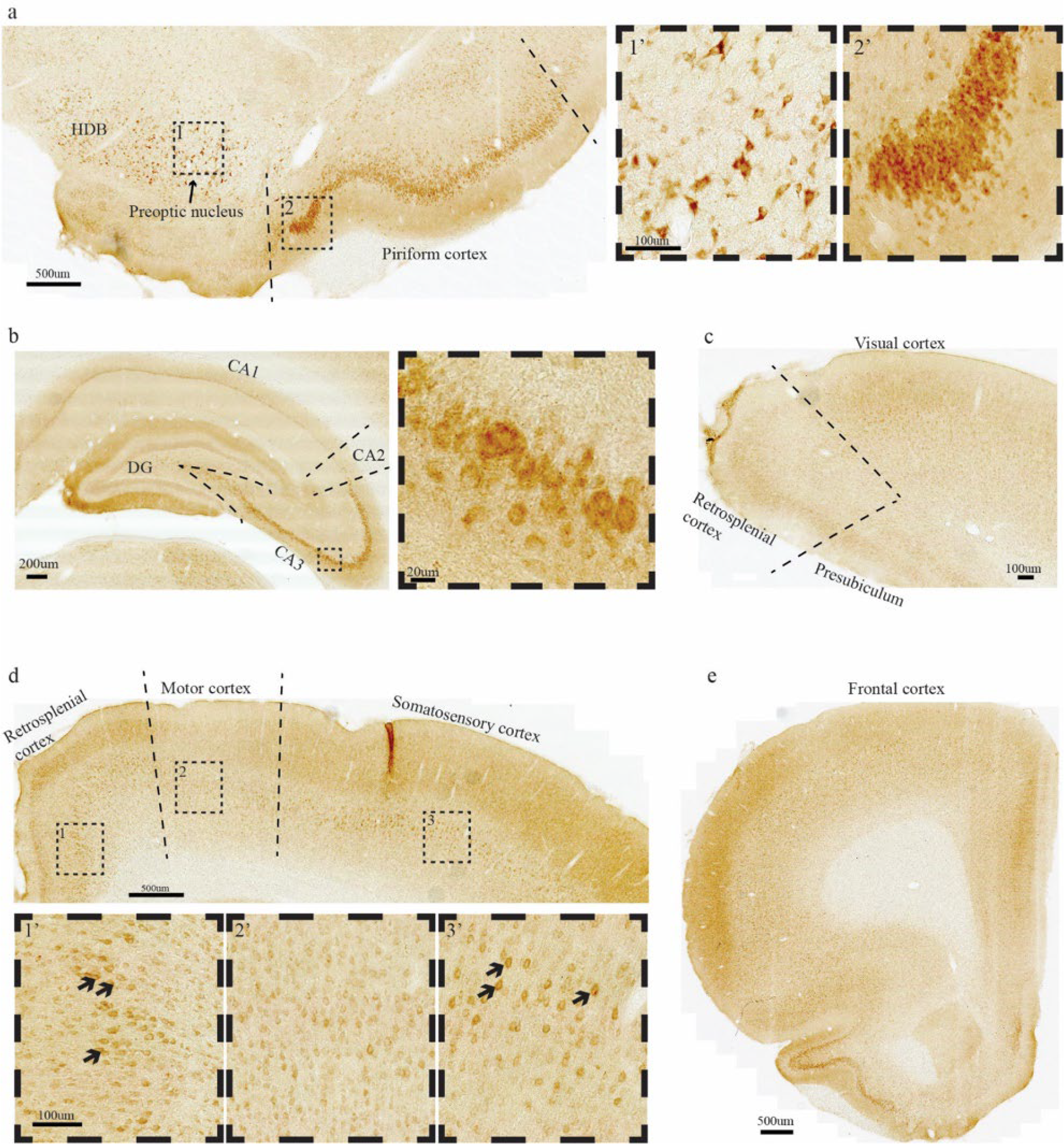
Moderate levels of Bnip3-expression is present in certain structures outside of the entorhinal cortex. This includes (a) neurons in the piriform cortex and the preoptic nucleus, (b) CA3 (also note the neuropil labeling in the outer 2/3ds of the molecular layer of DG, CA3 and CA2), (c) visual cortex, (d) retrosplenial-, motor-, and somatosensory cortex, and (e) frontal cortex. Scale bars are indicated in each figure, note that for the related insets, the leftmost scale bar applies to all.

Bnip3-expression in neurons of the dorsomedial part of the locus coeruleus (LC) is also high (Fig. 7 a, b).

**Figure 7.**
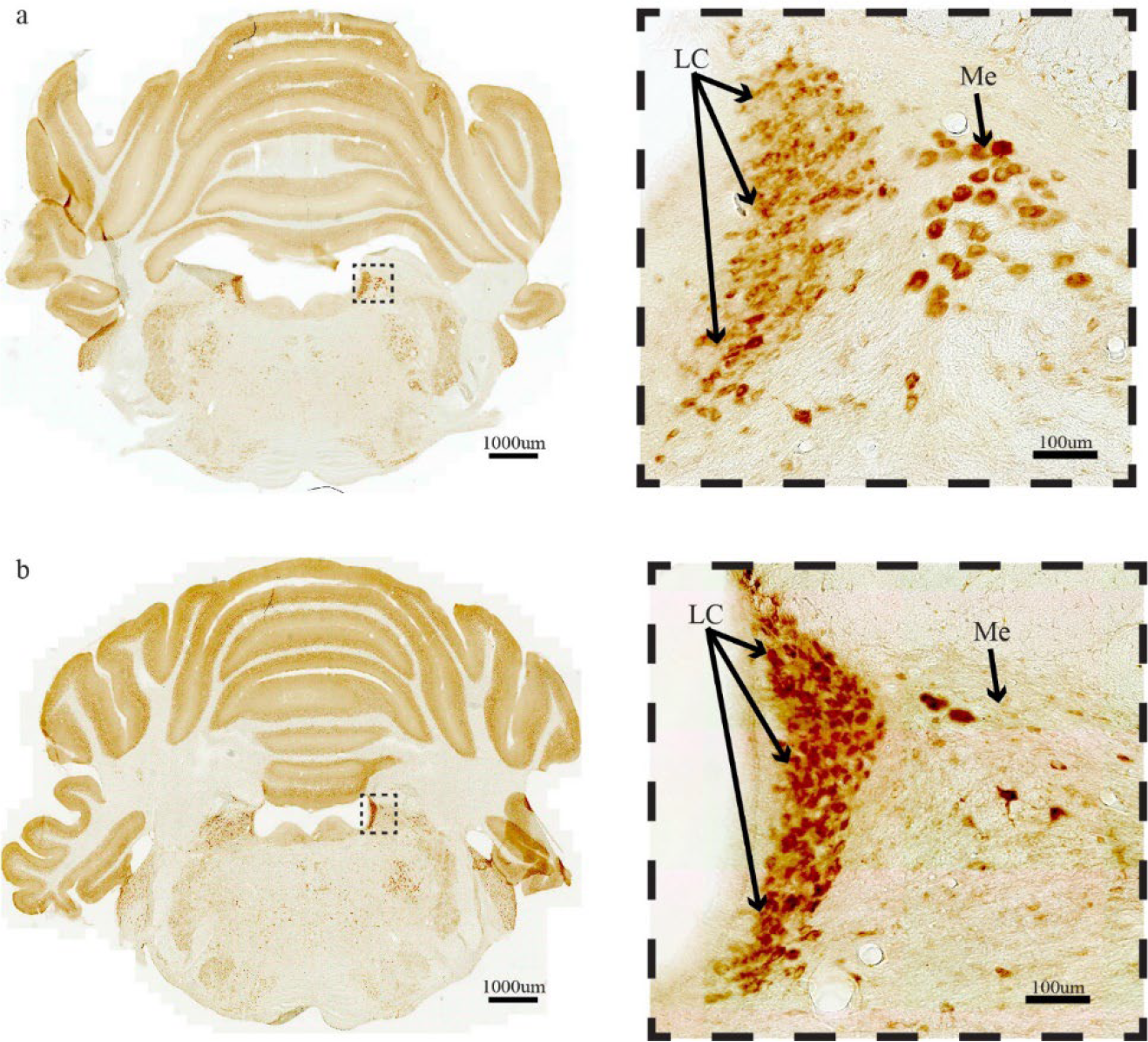
High levels of Bnip3-expression in neurons of the locus coeruleus (LC). (a) three month old rat, (b) 18 month old rat. Equally high levels are found in a subset of neurons belonging to the mesencephalic trigeminal nucleus (ME). Scale bars are indicated in each figure.

We confirmed that these Bnip3 high-expressing neurons are part of LC by double-labeling against tyrosine hydroxylase (Fig. 8). The high Bnip3 expression in these neurons suggests that they also have a higher than average metabolic rate, but it is beyond the scope of this paper to elaborate on the reasons for this.

**Figure 8.**
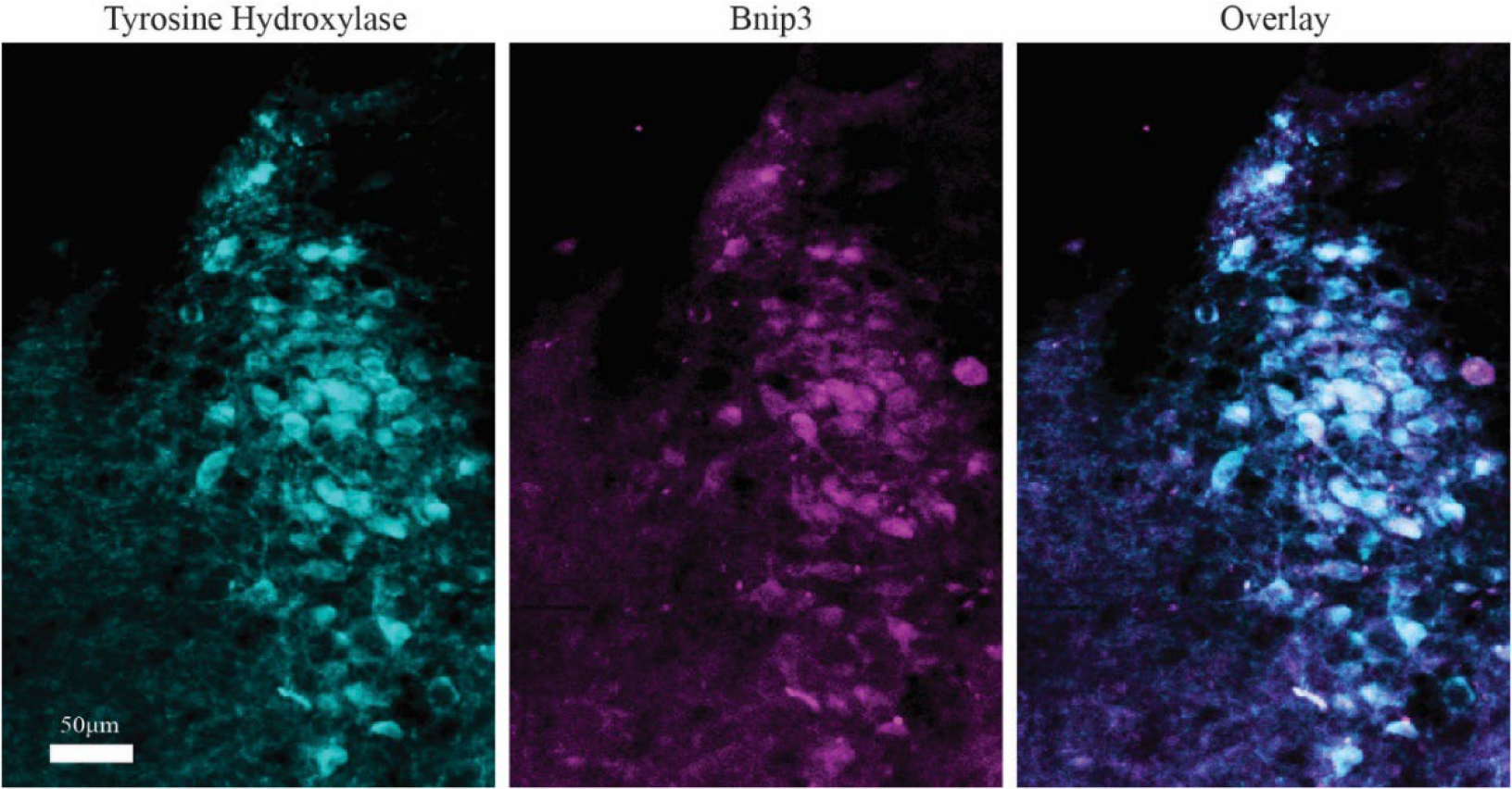
Bnip3-expression in neuron of LC co-localize with tyrosine hydroxylase. Scalebar in leftmost image applies to all.

## Discussion

Our results clearly support that, compared to most other neurons, Re+ ECLII-neurons generally have a particularly high energy requirement. And because the size of the standing pool of mitochondria is homeostatically regulated^33^ under normal conditions, enhanced Bnip3 expression also implicates enhanced mitogenesis. The observed expression gradient of Bnip3 shows that even though the metabolic rate in Re+ ECLII-neurons is generally high, it varies substantially in a systematic way. The systematic reduction in metabolic rate as one moves away from the rhinal fissure is in line with the distribution of the mitochondrial enzyme cytochrome oxidase in EC^34^.

We suggest that the reason why Re+ ECLII-neurons display the reelin and Bnip3 gradients is tied to the level of spatial and temporal detail by which they encode information about the external environment. In rodents, EC-neurons located close to the rhinal fissure are more strongly connected to the most dorsal part of the hippocampal formation, while successively more ventrally situated EC neurons are most strongly connected to correspondingly more ventral levels of the hippocampus^3,35^. Intriguingly, the level of detail by which spatial information about the external environment is encoded is highest at the dorsal extreme of the hippocampus (or the corresponding posterior extreme in humans) and gradually decreases as one moves ventrally (or anteriorly in humans) ^36-38^. Arguably, the more detailed the encoded environmental representation, the faster it will become outdated. Thus, to keep track of changes in the spatial environment, high-resolution representations are putatively generated and modified by the entorhinal-hippocampal system much more frequently than low-resolution ones. This conception is consistent with data on the only type of entorhinal neuron for which we have systematic data about the level of detail encoded for different populations, namely the grid cells in MEC, whose firing form a representation of space by tessellating the environment explored by the animal to form a hexagonal grid pattern^39^. Thus, grid cells with the most detailed grid pattern sit close to the rhinal fissure, while grid cells with increasingly less detailed grid patterns are located increasingly farther away from the rhinal fissure^40^. It is reasonable to expect that the need to update representations of the environment follows the scale under consideration. For example, several relevant details will change between the different times a rodent traverses a given stretch in its natural habitat, but the distal landmarks remain the same. On the assumption that levels of detail will prove to be encoded along the same gradient also for other, non-spatial representations of the environment, such as those of time or event sequences^41^, and objects^42^, we anticipate that the graded expression levels of reelin and Bnip3 will turn out to be directly related to the frequency with which it is necessary to update environmental representations in the broad sense. Since reelin is a canonical synaptogenic protein^12,13^ heavily involved in the remolding of synaptic contacts^43^, it makes at least perfect sense that its expression is highest in those neurons where the environmental representations are most frequently updated. And since this remolding must necessarily also be energetically demanding, this appears to explain why the Bnip3 expression is so closely tied to the reelin expression in Re+ ECLII neurons.

## Declarations

### Ethics approval

All procedures were approved by the Norwegian Animal Research Authority, and follow the European Convention for the Protection of Vertebrate Animals used for Experimental and Other Scientific Purposes. The Regional Committees for Medical and Health Research Ethics, Norway (permit 82685) approved the use of archival postmortem human brain sections.

### Consent for publication

All authors consent to publication of this manuscript.

### Availability of data and material

Raw data will be made available upon request.

### Authors contributions

Asgeir Kobro-Flatmoen conceived of and designed the study. Asgeir Kobro-Flatmoen and Raissa Lejneva collected the rat data, Asgeir Kobro-Flatmoen and Stig Omholt analysed the rat data. Asgeir Kobro-Flatmoen and Maria Jose Lagartos Donate collected and analysed the human data. Stig Omholt, Evandro Fei Fang and Asgeir Kobro-Flatmoen wrote the paper.

## Acknowledgements

We wish to thank Bruno Monterotti, Hanne Tegnander Soligard, Paulo Jorge Bettencourt Girão, and Grethe Mari Olsen for excellent technical assistance.

